# Limbic progesterone receptors regulate spatial memory

**DOI:** 10.1101/2022.08.09.503321

**Authors:** Suchitra Joshi, Cedric L Williams, Jaideep Kapur

## Abstract

Progesterone and its receptors (PRs) participate in mating and reproduction, but their role in spatial declarative memory, is not understood. Male and female mice express PRs in regions that support spatial memory: the hippocampus and entorhinal cortex. PRs were predominantly expressed in excitatory neurons in *Pgr*-Cre mice injected with AAV-delivered flexed tdTomato. Furthermore, segesterone, a specific PR agonist, activated neurons in the entorhinal cortex (EC) and the hippocampus. We assessed the PR function in these spatial memory circuit neurons by examining the performance of mice lacking this receptor (PRKO) in novel object recognition, object placement, and Y-maze alternation tasks. In the recognition test, wild-type littermates spent significantly more time exploring the new object than male PRKO mice. The EC-specific deletion of PRs was sufficient to induce a deficit in detecting familiar versus never experienced or new objects. We confirmed deficits in spatial memory of PRKO mice by testing them on the Y-maze forced alternation task. In contrast to spatial tasks, PR removal did not alter the response to fear conditioning. PRs also support spatial memory in female mice. These studies provide novel insights into the role of PR in facilitating spatial, declarative memory in males and females, which may help with finding reproductive partners.

**SIGNIFICANCE STATEMENT:** Brain progesterone receptors play an essential role in facilitating mating and reproductive behaviors, but their role in spatial memory and, therefore, mate-finding is not described. Principal, excitatory neurons of the entorhinal cortex and hippocampus express progesterone receptors. These receptors facilitate spatial memory in male and female mice, which may enhance mate-finding reproductive function and food foraging.

## INTRODUCTION

Progesterone, a canonical female reproductive hormone, is also present in males. Men’s progesterone levels are similar to postmenopausal women’s (Oettel and Mukhopadhyay, 2004). The brain synthesizes progesterone and other neurosteroids from cholesterol; indeed, brain pregnanolone levels are thirty times higher than serum levels (Baulieu, 1998; Schumacher et al., 2014; Zhu et al., 2017). Male mouse brain progesterone levels are comparable to those in females in the diestrus stage (Zhu et al., 2017), and remain stable. Progesterone suppresses sexual behavior in males by antagonizing androgen actions (Witt et al., 1994). Progesterone receptors (PRs) are critical for this regulation of reproductive behavior since their genetic ablation (PR knockout) enhances male sexual behavior (Schneider et al., 2005). PR deletion also reduces aggression toward other males or newborn pups (Schneider et al., 2003). Thus, these progesterone-PR actions in males are opposite of those in females. However, other PR functions are unknown.

Through conversion to its metabolite allopregnanolone, progesterone rapidly enhances memory in experimental animals (Frye et al., 2007; Frye and Walf, 2008; Lewis et al., 2008; Frye and Walf, 2010; Frye et al., 2021). These effects do not seem to be impacted by PR deletion (Frye and Walf, 2010; Frye et al., 2021). This is not a surprising finding since the animals were tested four hours after progesterone treatment; this time may be inadequate to uncover effects of PRs, which alter gene expression and occur relatively slowly (Brinton et al., 2008; Mani and Oyola, 2012). In contrast, PR blockade during development appears to alter novel object recognition performance when tested later in life in adult males (Newell et al., 2018). Unfortunately, no studies have investigated the role of PRs in regulating cognition in adult rodents.

To this end, the performance of wildtype and PR knockout (PRKO) were examined on spatial, short- and long-term tests of memory with the use of a novel object recognition, object place, and Y-maze tasks. The involvement of PR’s in regulating emotional memory was assessed in both groups with a standard contextual fear conditioning test. Anatomical specificity of PR effects was evaluated by measuring PRs expression in the entrorhinal cortex (EC) and hippocampi using qRT-PCR, and then localized their neuronal expression using mice expressing a Cre recombinase under the regulation of *Pgr* promoter. We measured PR activation-induced neuronal activity using TRAP2 mice. In summary, the collective experiments were conducted to reveal the distribution of EC and hippocampal PR’s that regulate spatial and emotional memory.

## MATERIALS AND METHODS

### Animals

Mice with a floxed first exon of *Pgr* (PRCE) were a kind gift from Dr. M. Luisa Iruela-Arispe (University of California, Los Angeles, CA) (Hashimoto-Partyka et al., 2006; Joshi et al., 2018). The mice were crossed with nestin-Cre mice (B6.Cg-Tg(Nes-Cre)1Kln/J, The Jackson Laboratory # 003771) to generate animals with brain-specific PR deletion. The colony was maintained by breeding the male mice with brain-specific deletion of PRs (denoted as PRKO here onwards) with PRCE homozygous females. Mice expressing Cre recombinase under the control of Pgr (B6.129S(Cg)-Pgrtm1.1(Cre)Shah/AndJ, The Jackson Laboratory #017915) were also used. An advanced version of TRAP mice described in our earlier study was also used (Allen et al., 2017). All animals had ad libitum access to food and water and were maintained on a 12-hour light and 12-hour dark cycle (lights on at 6 AM, lights off at 6 PM). All experiments were performed with 50-70 day-old animals.

### Behavior testing

The animals were handled for three days, five minutes each before behavioral testing. Some animals were treated with RU 486 (10 mg/kg, subcutaneous, sc; n=9 vehicle-treated and 7 RU-486-treated), a PR antagonist that should produce similar effects as the PRKO, or vehicle (20% β-hydroxycyclodextrin) for a week before behavioral testing. A cohort of animals was also treated with progesterone (10 mg/kg/day, sc) for two days before testing (Joshi et al., 2018), to determine if excitatory actions of PRs in the EC and hippocampus enhance memory (n= 9 vehicle-treated and 10 progesterone-treated WT and 6 vehicle-treated and 7 progesterone-treated PRKO mice).

#### Novel object recognition test

Non-spatial working memory was evaluated using a novel object srecognition task (Leger et al., 2013). Briefly, the animals were habituated to the arena (45 × 45 × 40 cm, Harvard Apparatus) for 10 min for two days (habituation) before the experiment (Fig. 3A). On the experiment day, a total of 14 WT and 15 PRKO were exposed to the arena (familiarization phase) with two identical objects placed at the opposite corners and allowed to explore for 5 min. The animals were returned to their home cages for a delay of either one or 8 hours and then re-exposed to the arena (testing phase) for 5 min. During the recognition test, one of the original objects was replaced with a novel object that was similar in size but differed in both color and shape. A video recorder and AnyMaze software (Stoelting) were used to quantify exploration behavior during the familiarization and testing phases. The three measures of exploration included 1) time spent in the target zone (i.e. 2 cm surrounding the objects), 2) total interaction with the objects and 3) percent time spent with the novel object [i.e. time with novel object)/ (total time spent with the two objects)] X 100]. The animals were excluded from the experiment if the total object exploration time was less than 20 sec during the familiarization phase.

#### Object Place assay

The procedure to assess spatial memory discrimination was similar to the novel object recognition test described above. The only deviation occurred during the testing phase, where the location of one of the original objects explored during the familiarization phase was moved to a new location in the testing phase (Fig. 4A). Thus, the novel experience during this test involved the new location of a familiar object. Eight WT and PRKO mice each were used for these studies.

#### Y maze forced alteration assay

The animals were handled three days before the experiment. The animals were exposed to the Y maze apparatus for 10 min with one of the two arms blocked (Fig. 4D). After the first exposure, the animals were returned to the home cage for 30 minutes and re-exposed to the apparatus for another 10 minutes with both arms open. The number of entries into novel vs. familiar arms and the percentage of time spent in the novel vs. familiar arms were compared among animals. If previous experience with a familiar arm is encoded into memory, animals tend to spend less time re-exploring the remembered context and show an increase in exploration of the novel addition to the testing arena. Six animals of each genotype were used in these studies.

#### Contextual fear conditioning

A chamber (27 × 21 × 21 cm; Med Associates Inc) equipped with an infrared camera at the front for recording the behavior was used for contextual fear conditioning. The chamber floor consisted of stainless-steel rods placed 6 mm apart. A standalone aversive stimulator applied foot shock via the floor rods. An audio generator– connected speaker at the chamber top provided an auditory signal, with a background noise of 60 dB supplied by a fan at the side of the chamber. The chamber was cleaned with 70% alcohol in between animals and wiped with 1% acetic acid before the animal exposure. For conditioning, the animals were placed in the chamber and allowed to explore for 3 min before the application of five 2-sec foot shocks paired with a tone (90 Db, 5000 Hz, 20 sec). The shock (0.5 mA for 2 sec) was contiguous with the last 2 sec of the tone. Each tone-shock stimulus was separated by 60 sec. The animals were removed from the chamber 2 minutes after the last shock and returned to the home cage. Video Freeze software was used to track the animal and to apply tone and foot shock. Six WT and 7 PRKO mice were used in these studies.

We tested context recall 24 hours after conditioning. After cleaning the chamber with 1% acetic acid, the animal was placed in it for 8 minutes without tone or shock. Freezing was defined as immobility for 2 sec, and the freezing time was expressed as a percent fraction of total time in the chamber.

### qRT-PCR

Hippocampus and EC were isolated from male and gonadally-intact female mice in the estrus stage of the cycle. Uterine tissue of female mice in the estrus stage was used as a control for normalization. Total RNA isolation and cDNA synthesis was performed as described before (Joshi et al., 2018). The qPCR assay was performed using sensiFAST SYBR and fluorescein kits (BioLine) and primers used in prior studies(Turgeon and Waring, 2006; Stephens et al., 2011); PGR-F: TCTACCCGCCATACCTTAACT and PGR-R: GTGACAGCCAGATGCTTCAT and GAPDH-F: ACAGTCCATGCCATCACTGCC and GAPDH-R: GCCTGCTTCACCACCTTCTTG. The gene expression was determined using the 2^-ΔΔCt^ method, with the expression in the uterine tissue taken as a control for normalization (Livak and Schmittgen, 2001). The PCR cycle consisted of 95 °C for 5 min, and 40 cycles of 95 °C for 15 sec followed by 60 °C for 15 sec. Negative controls included reactions without RT. The validation of the qRT-PCR assay is provided in supplementary figure 1. The PCR amplicon was sequenced to confirm the amplification of the correct target. To validate the 2^-ΔΔCt^ method, Ct values for PR and GAPDH were determined for cDNA dilutions ranging from 1:10 to 1:100000. The log cDNA dilution was plotted against ΔCt; the absolute slope of the line was 0.03, indicating that the efficiency of amplification of PR and GAPDH was similar over a wide range of copy numbers (Livak and Schmittgen, 2001). The Cvs. log copy number plots were comparable for the PR and GAPDH PCRs (Fig. 1C), and the amplification product was absent in the PRKO animals (Fig. 1A, B).

**Figure 1:**
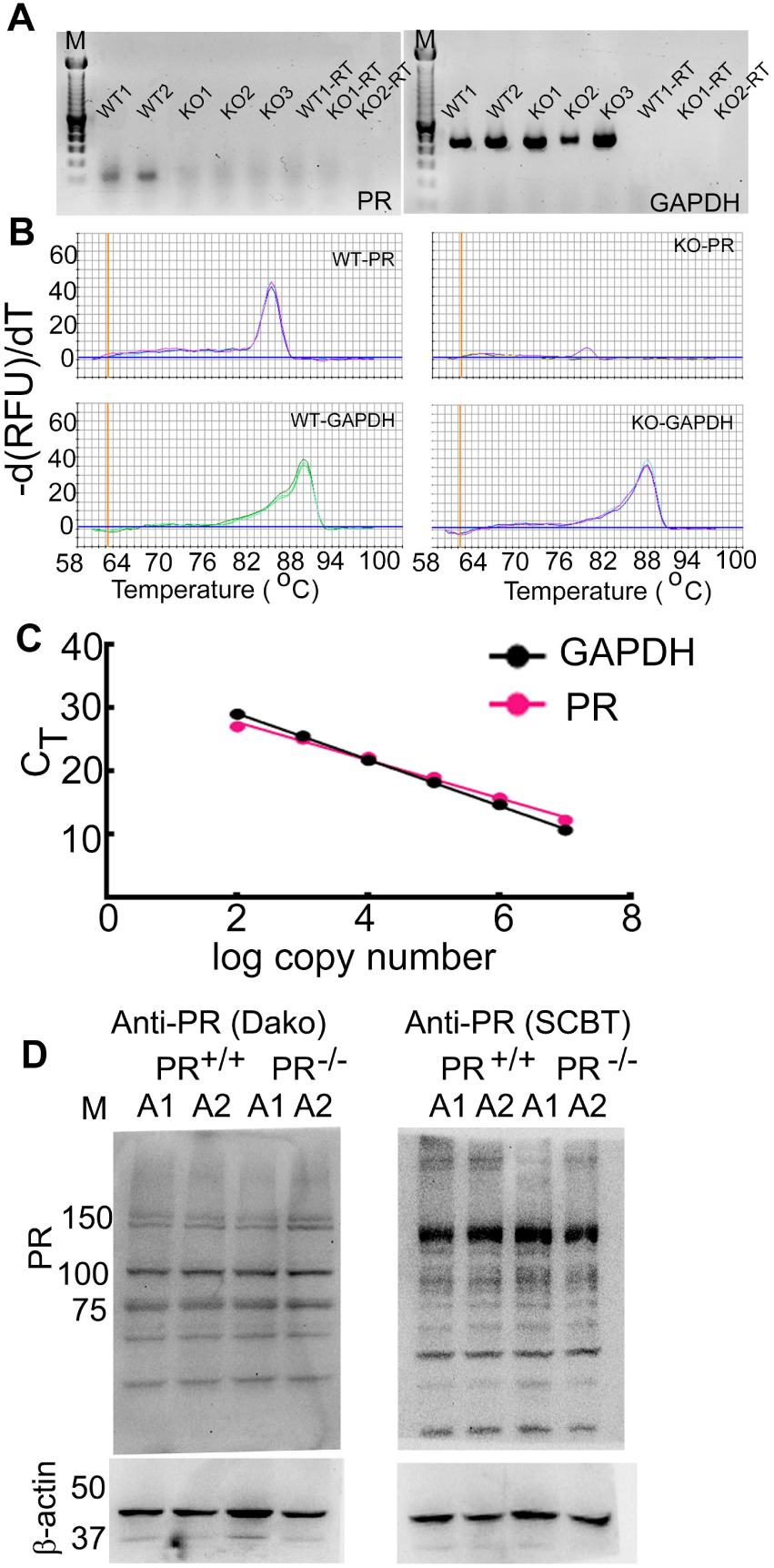
Validation of the qRT-PCR assay. **(A)** A PCR was performed using primers amplifying a fragment of PR and GAPDH using cDNA as a template. The PCR reaction generated a single band of the expected size in the WT animals; an amplification product was not seen in the PRKO animals. **(B)** Melting curves of PR and GAPDH qPCR from representative PRKO and WT mice. **(C)** Efficacy of amplification of PR and GAPDH primers. **(D)** Western blots of brain proteins using two anti-PR antibodies. The expression of β-actin was used as a loading control. Molecular weight markers are shown in lane M. Please note that the antibodies reacted with multiple proteins suggestive of lower specificity.

### TRAPing of active neurons

TRAP2 male mice were housed four per cage. Segesterone (10 mg/kg, subcutaneous, sc, a single dose) or vehicle (20% β-hydroxycyclodextrin) were administered to the mice, and two days later, a synthetic estrogen receptor (ER) ligand 4-hydroxytamoxifen (4-OHT, 50 mg/kg, sc) was administered. The cage was kept in a quiet area, and the mice were not handled on that day before the 4-OHT injection to avoid neuronal activation due to other stimuli. Seven days after 4-OHT administration, the animals were transcardially perfused with a solution containing 4% paraformaldehyde and 1% acrylamide and post-fixed in the same overnight. As described before, two hundred micron-thick horizontal brain sections were processed for passive tissue clearing (Dabrowska et al., 2019). The expression of neuronal marker protein NeuN (anti-NeuN, 1:200, MAB377, Millipore) was determined to identify neuronal tdTomato expression.

### Image acquisition and analysis

The tissue was imaged using a Nikon Eclipse Ti-U microscope equipped with a Nikon confocal C2 scanner, DU3 High Sensitivity Detector System, and Nikon LUN4 4 Line Solid State Laser System under 10X 0.45 numerical aperture lens with 512 × 512 frame size. Ten µm optical sections were acquired, and images were tiled and stitched using NIS-Elements software. The images were processed and analyzed using Imaris 8.3.1 software (Bitplane Scientific, Zurich, Switzerland). The regions of interest were marked using Paxinos Atlas (Paxinos G, 1986).

### Measurement of progesterone levels

Combined EC and Hp tissue from WT and PRKO male mice was isolated, snap-frozen, and stored at −80°C until use. The steroid levels were measured using the DetectX progesterone enzyme immunoassay kit (Arbor Assays, Ann Arbor, MI) with a 50 to 3200 pg/mL detection range. The steroids were extracted from the tissue with acetonitrile using a steroid tissue extraction protocol (Arbor Assays). The steroids were suspended in 50 μl ethanol, and 10 μl of the suspension was used for the assay. All the samples were run in duplicate.

### Ovariectomy

In female mice, we removed ovaries bilaterally under isofluorane anesthesia (Shiono et al., 2021) and allowed them to recover for 10-12 days. The circulating progesterone and estrogen levels were measured in the ovariectomized animals using ELISA (Shiono et al., 2021). The circulating estrogen (<3 pg/ml, n= 4) and progesterone (0.38 ± 0.42 ng/ml, n= 4) levels were negligible in these animals.

### PR deletion in the EC

AAV9 expressing GFP-Cre under the control of CamKII promoter (pENN.AAV.CamKII.HI.GFP-Cre.WPRE.SV40, Addgene #105551) was used to delete the expression of PRs in the EC of adult PRCE males. Control mice were injected with AAV9 expressing GFP under the control of CamKII promoter (pENN.AAV.CamKII0.4.eGFP.WPRE.rBG, Addgene # 105541). The injections were performed bilaterally (Adotevi et al., 2019). The stereotaxic coordinates used for viral injections were anteroposterior [AP] −4.6; mediolateral [ML], −/+3.0; dorsoventral [DV], −3.7 and-3.0 from dura for central and medial EC and AP −3.7, ML −/+ 4.2, and DV −3.5 and −3.0 from dura for lateral EC. A Hamilton Company (Reno, NV) syringe (Hamilton 7000 Glass, one μl, 0.3302mm) was loaded with virus solution and mounted in the peristaltic pump holder (Harvard Apparatus, Holliston, MA; P-1500), 200 nl of the virus was injected at each site at a flow rate of 200 nl/min. The experiments were performed two weeks after viral injection. At the end of the experiment, viral transduction of the EC cortical neurons was confirmed by evaluating GFP expression in the brain slices.

AAV9 expressing a flex tdTomato (pAAV-FLEX-tdTomato, Addgene # 28306) was used to evaluate PR expression in the EC-hippocampal network in the Pgr-Cre mice. A 100 nl of the virus was injected unilaterally in the medial/central EC and lateral EC (please see the stereotaxic coordinates above), in the DG (AP −3.6, ML 2.5, DV −2.3), and the CA1 (AP −3.0, ML 3.5, DV −3.5); the animals were perfused 14 days after virus injection, 40 μm-thick sections were prepared and co-labeled with NeuN as described above. The sections were also immunolabeled with anti-CamKII (1:500, ab34073, Abcam) to determine whether the principal neurons expressed PRs. The sections were labeled with anti-GluA2 (1:1000, clone 6C4, Millipore), anti-somatostatin (1:500, ab108456, Abcam), and anti-parvalbumin (1:500, ab427, Abcam) antibodies using standard protocols (Sun et al., 2007) to evaluate PR expression in the interneurons and hilar mossy cells.

### Statistical analysis

Graphpad Prism 9 was used to perform statistical comparisons. All the values are expressed as mean ± standard deviation. The differences were considered significant when the p-value was less than 0.05. The normal distribution of the data was determined using the Kolmogorov-Smirnov test. The normally distributed data were compared using the student’s t-test or ANOVA followed by šídák’s multiple comparisons test.

## RESULTS

### EC and hippocampus of adult male mice express PRs

Previous studies described PRs expression in the hypothalamic nuclei of male mice but not in the limbic regions. Furthermore, no quantitative comparison of the PR expression in the EC and hippocampus of male and female animals is available. We evaluated PR mRNA expression in the EC and hippocampal tissue using a qRT-PCR assay. The PR mRNA was present in the EC and hippocampi of male mice (Fig. 2E), and their levels were comparable to those in the female animals in the estrus stage of the cycle. The EC PR mRNA levels were similar to those in the uterus (standard), and twice that in the hippocampus.

**Figure 2:**
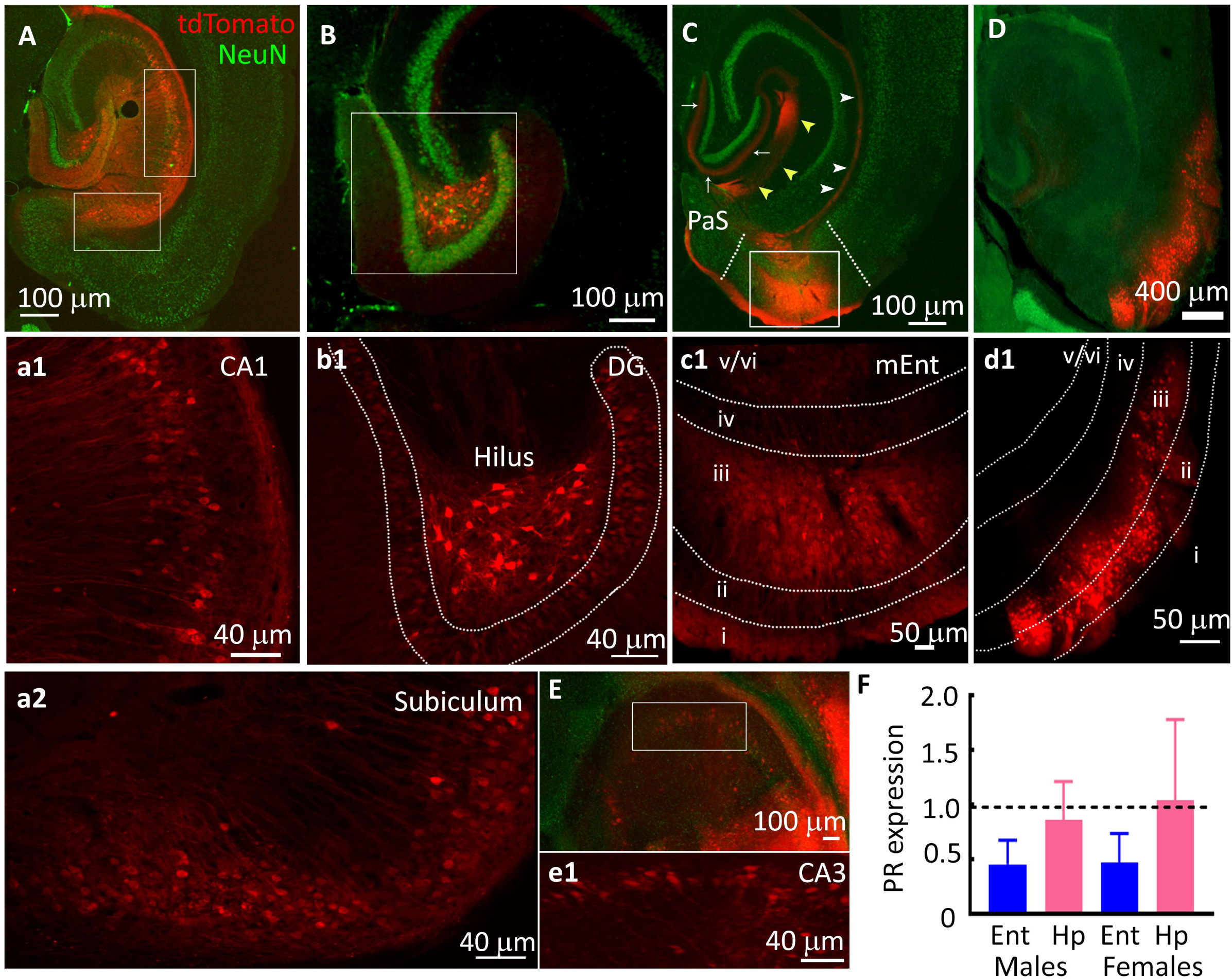
PR expression in the EC and hippocampus of adult male mice. **(A-E)** Images from representative Pgr-Cre animals injected with AAV9 expressing a flexed-tdTomato. The AAVs were injected into the hippocampus and entorhinal cortex (EC). The sections were counterstained with a neuronal marker protein NeuN (Green) to label neurons. The tdTomato labeled terminals of the perforant path (white arrows), temporoammonic pathway (white arrowheads), and temporoammonic Alvear path fibers and stratum lacunosum moleculare (yellow arrowheads) are also marked in 1C. (**a1, a2, b1-e1**) Magnified boxed regions from A-E showing neuronal tdTomato expression. **(F)** PR mRNA expression in the EC and hippocampus (Hp) of male mice, n= 7 for EC and 5 for Hp. The PR mRNA expression (1.12± 0.29, n=7) in the uterine tissue isolated from a female in the estrus stage of the cycle was used for normalization (dotted black line). The expression of PR mRNA in this tissue isolated from female mice in the estrus stage is shown for comparison (n=6 each).

We then studied the cellular distribution of PRs in the EC and hippocampus. None of the commercially available anti-PR antibodies had the specificity necessary for immunohistochemical detection (Fig. 1D). Therefore, we used mice expressing a Cre recombinase under the control of the *Pgr* promoter to define the cellular distribution of PRs. An AAV9 (with flexed tdTomato) was injected in the regions of interest in these mice to label the PR-expressing cells with tdTomato fluorescence.

The tdTomato expression was evident in the CA1 and subicular neurons (Fig. 2A, 1a1, 1a2). In the CA1 region, the labeled neurons were present in the stratum pyramidale and stratum oriens. The DGC tdTomato expression was weaker than in the CA1 and subiculum (Fig 2B, 1b1). The tdTomato expression in the CA3 region was also sparse (Fig. 2E). Several neurons in the dentate hilus also expressed tdTomato (Fig. 2B, 1b1). The neurons of all the three EC subdivisions, medial, central, and lateral, also expressed tdTomato (Fig. 2C, 1D, 1c1, 1d1). sWithin the central EC, the tdTomato-expressing cells were commonly present in layer III. On the other hand, the mEC and lEC tdTomato expression was spread in layers II/III. The layer II/III EC neurons project to the DGCs and CA1 neurons through the perforant pathway (PP) and temporo-ammonic pathway (TA), respectively; the terminals of the labeled EC neurons marked the molecular layer of the dentate gyrus (Fig. 1C). There was tdTomato fluorescence in the temporo-ammonic Alvear path fibers and stratum lacunosum moleculare of the CA1 hippocampus (Fig. 2C).

PRs were preferentially expressed in the principal neurons. We evaluated tdTomato colocalization with CamKII protein, a marker of glutamatergic neurons. Indeed, the tdTomato labeled DGCs, CA1, subicular, and EC neurons colocalized with CamKII (Fig. 3D, 2G, 2J, and 2M). We evaluated tdTomato fluorescence colocalization with parvalbumin (PV) and somatostatin (Som) peptides to confirm the absence of PR expression in the interneurons. None of the tdTomato-positive neurons in the DG cell layer, CA1, subiculum, and EC colocalized with PV (Fig. 3B, 2E, 2H, 2K, 2N). The tdTomato expression did not colocalize with the Som immunoreactivity. Only a few neurons in the hippocampus and EC expressed Som (Fig. 3C, 2F, 2I, 2L, 2O).

**Figure 3:**
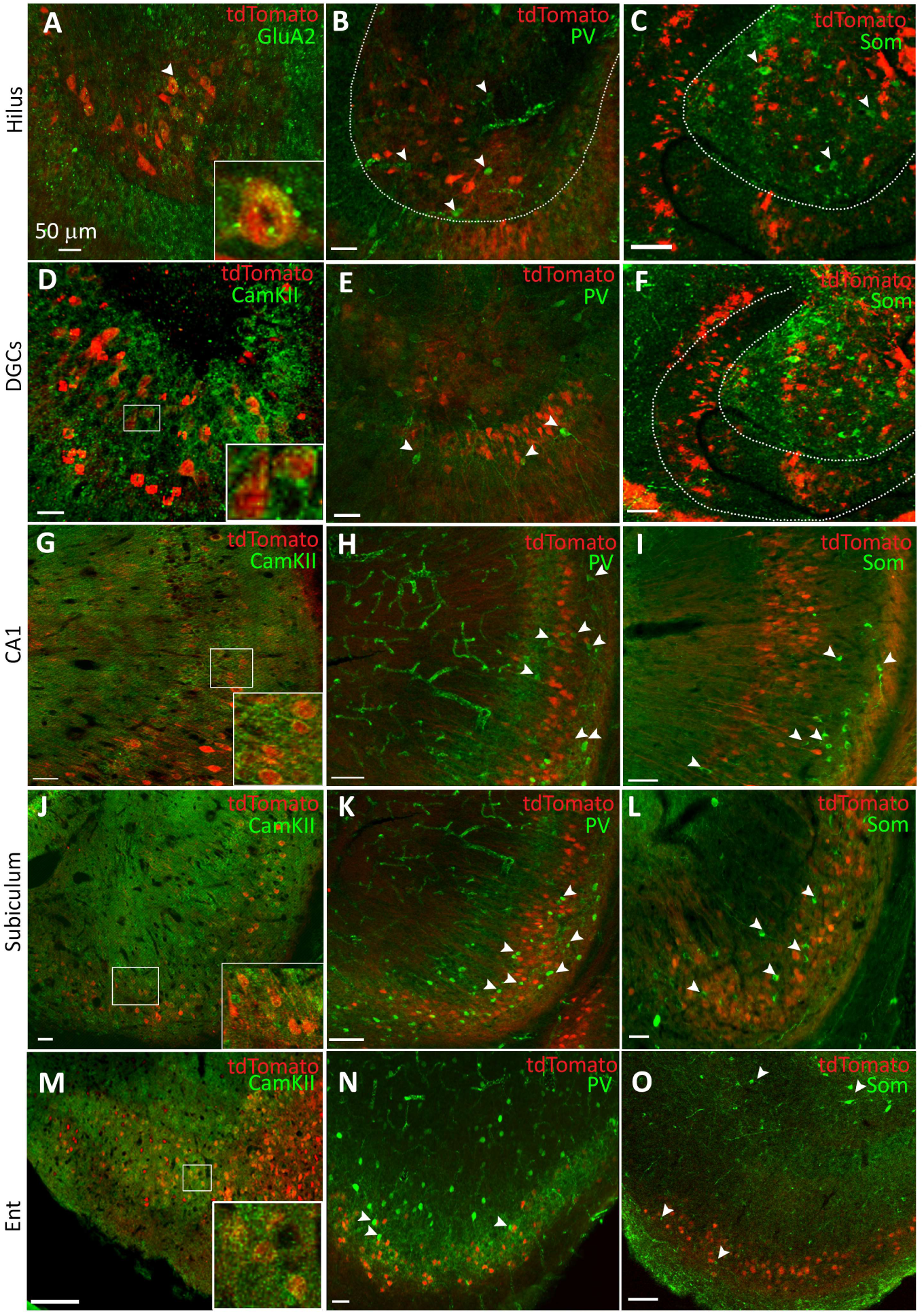
PR expression in the principal neurons vs interneurons. **(A, B, C)** Representative images of the dentate hilus of Pgr-Cre mice injected with AAV expressing flex tdTomato. The images show colocalization of tdTomato expression with that of the immunoreactivity of the GluA2 subunits of AMPA receptors, parvalbumin (PV), and somatostatin (Som) neuropeptides respectively. The expression of the GluA2 subunit marked the hilar mossy cells, whereas, parvalbumin and somatostatin labeled two types of interneurons. The scale bars in these and all other images in this figure correspond to 50 μm. The hilar neuron marked by an arrow is magnified in the inset in A to show punctate GluA2 subunit immunoreactivity in a tdTomato-expressing neuron. The arrows in B and C mark the parvalbumin and somatostatin-expressing neurons respectively, none of which colocalized with tdTomato fluorescence. **(D, E, F)** Representative images showing colocalization of tdTomato with that of CamKII, PV, and Som respectively in the DGCs. The inset in D corresponds to the boxed region in D and shows perisomatic CamKII immunoreactivity in the tdTomato-positive DGCs. The arrows in E and F point to the interneurons. **(G, H, I)** Colocalization of tdTomato and CamKII, PV, and Som in the CA1. The boxed region in G is magnified in the inset, arrows in H and I point the interneurons. **(J, K, L)** Colocalization of tdTomato with CamKII, PV, and Som expression respectively. The boxed region in J is magnified as an inset, arrows in K and L point to the interneurons. **(M, N, O)** Colocalization of tdTomato in the medial EC with CamKII, PV, and Som respectively. Please note that som labeling in the superficial layers of EC was sparse, but som-positive neurons were present in the deeper layers (marked by arrows). Also please note that the number of som-expressing neurons seems to be substantially lower than those expressing PV.

There were tdTomato expressing cells in the dentate hilus, a region populated by inhibitory GABAergic interneurons and excitatory mossy cells (Scharfman, 2016). The expression of GluA2/3 subunits of AMPA receptors distinguishes mossy cells from interneurons (Leranth et al., 1996). Thus, to evaluate whether the tdTomato-expressing hilar neurons were mossy cells, we determined their colocalization with the GluA2 subunit protein. Many tdTomato-expressing hilar interneurons also expressed punctate and perisomatic GluA2 immunoreactivity (Fig. 3A). We tested whether the hilar tdTomato-positive neurons expressed PV or Som), the neuropeptides expressed in hilar inhibitory interneurons. As expected, hilar neurons expressed PV or Som, but none of these neurons expressed tdTomato (Fig. 3B, 2C). Thus, PRs were expressed primarily in the mossy cells and not on PV or Som-expressing inhibitory interneurons. Thus PRs were primarily in the excitatory neurons expressed. We then tested the effect of activating PRs in EC and the hippocampus.

### PRs activate neurons in the EC cortex and hippocampi of male mice

PRs increase EC neurons’ excitability in female rats (Shiono et al., 2021). We, therefore, expected that a similar PR effect would increase neuronal activity specifically in PR-expressing neurons but not in the regions lacking PRs. We gave segesterone or saline to the activity-reporter targeted recombination in active populations-2 (TRAP2) mice in their home cage(Allen et al., 2017) to test the effect of activating PRs. The active neurons were TRAPed two days later by administration of 4-hydroxytamoxifen(4OHT).

Segesterone treatment enhanced EC neuronal activity. There were tdTomato-expressing neurons in the EC of vehicle- and Segesterone-treated animals (Fig. 4B, 3B); however, in the Segesterone-treated animals, there were many more labeled neurons than in the vehicle-treated controls. The labeled neurons were present in all the subfields of the EC: central, medial, and lateral. Within the medial EC, the labeled neurons were concentrated in layers II/III and V, whereas, in the lateral EC, they were present in all the layers. Like in EC, Segesterone treatment also activated hippocampal neurons. Neurons in the DG, CA1 and subiculum expressed tdTomato (Fig. 4). Although PR-expressing neurons were present in the dentate hilus, the tdTomato-expressing neurons were sparse in this region (Fig. 4C, 3D). We counted all the labeled neurons in both groups within EC and CA1+subicular regions and DGCs. Segesterone treatment activated twice as many EC neurons (Fig. 4I), CA1, and subicular neurons as the vehicle-treated animals (Fig. 4J). The number of active DGCs in Segesterone-treated animals was thrice that of the vehicle-treated animals (Fig. 4K). Segesterone treatment did not activate lateral septum or collicular neurons. The number of TRAPed neurons in the lateral septum and colliculus was similar in the segesterone and saline-treated animals (Fig. 4E-3H), suggesting a specific segesterone effect. Thus, segesterone activated s PR-expressing neurons or their downstream targets.

**Figure 4:**
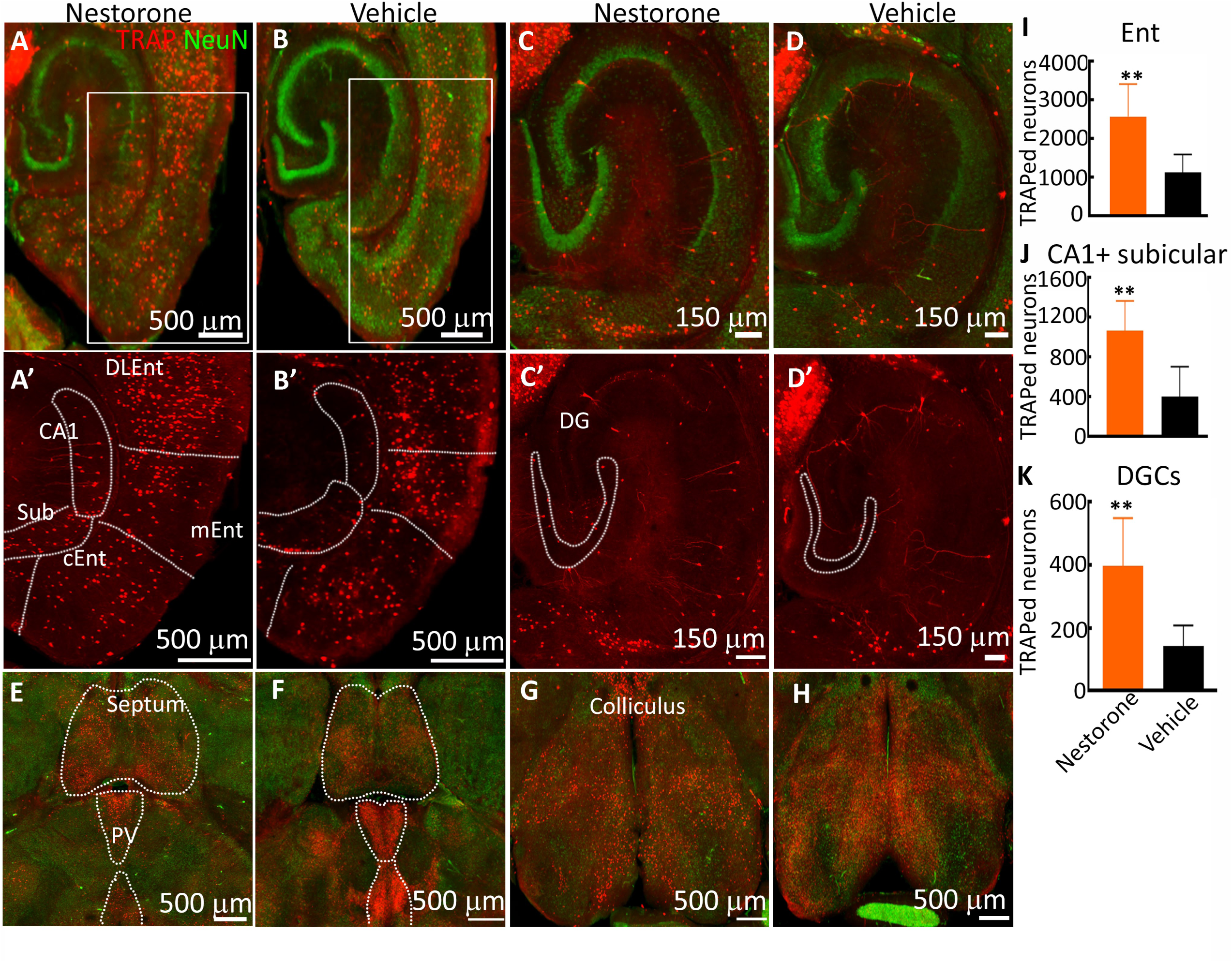
The effect of PR activation on the number of active neurons in the EC and hippocampus of adult male mice. **(A, B)** Representative images from segesterone-(10 mg/kg, subcutaneous) and vehicle-(20% β-hydroxycyclodextrin) treated male mice respectively. 4OHT was administered 2 days after the treatment in home cages to TRAP the active neurons (red). The green fluorescence corresponds to the expression of neuronal marker protein NeuN. **(A’, B’)** Magnified boxed areas from A and B respectively. The CA1, subiculum (sub), central (cEnt), medial (mEnt) and dorsolateral (dlEnt) EC regions are marked. **(C, D)** Representative images from segesterone- and vehicle-treated mice show active DGCs, the NeuN immunoreactivity is shown in green. **(C’, D’)** Images showing the TRAPed DGCs in the segesterone- and vehicle-treated animals. **(E, G)** and **(F, H)** Images showing the TRAPed neurons in the septum, paraventricular thalamic nucleus, and colliculus of segesterone- and vehicle-treated animals respectively. **(I, J, K)** The number of TRAPed neurons in the entorhinal cortex, combined CA1 and subiculum, and DGCs in segesterone- and vehicle-treated animals, n=5 each, ** p=0.0098 for EC (t=3.370, df=8), p=0.008 for CA1 and subiculum (t=3.508, df=8), and p=0.0094 for DGCs (t=3.394, df=8), student’s t-test.

### Blockade of PRs impacted object recognition in male mice

We further tested the role of PRs in EC and the hippocampus, which critically regulate spatial memory (Winters et al., 2008; Broadbent et al., 2009; Wilson et al., 2013b; Chao et al., 2016). Using an object recognition task, we compared declarative memory in male mice lacking PRs with WT controls. The animals were habituated and familiarized with two identical objects (Fig. 5A), and then tested after 8 hours in their home cages. We replaced one of the original objects with a novel object during the test. The WT animals explored the novel object longer than the familiar one. PRKO mice did not prefer the novel object (Fig. 5B) and spent less time with it than WT mice (Fig. 5C, p=0.029, t=2.499, df=11, student’s t-test). Next, we evaluated whether a shorter interval of 1 hour rather than 8 hours could alleviate the deficit in the PRKO mice. However, these mice were deficient even when tested an hour after the familiarization (Fig. 5D, p=0.0006, t=4.291, df=15, student’s t-test).

**Figure 5:**
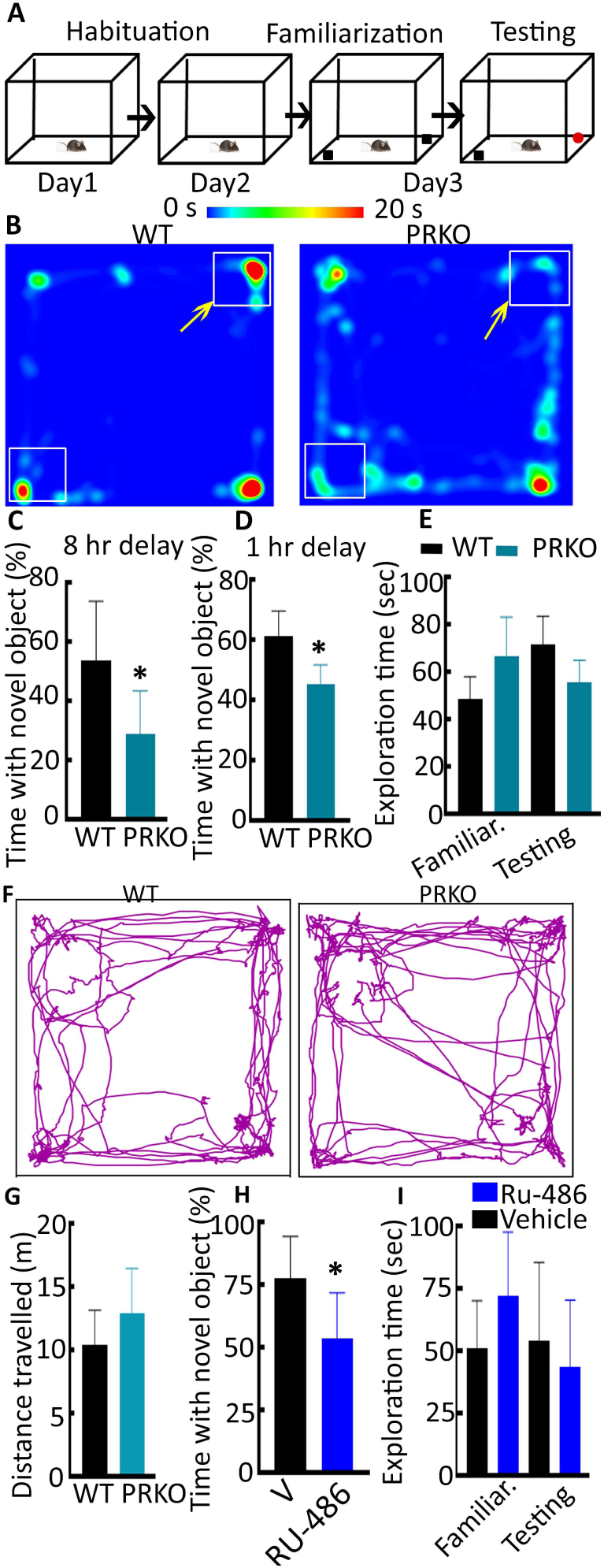
The effect of PR deletion on novel object recognition in adult male mice. **(A)** A schematic showing habituation familiarization, and testing of animals in a novel object recognition test. The assay was performed as described in the methods section. **(B)** A heat map of the time spent by representative PR knockout (PRKO) and littermate wildtype (WT) mice during the testing phase was performed 8 hours after the familiarization. The yellow arrows denote the position of novel objects. **(C)** The average and SD of the percent novel object exploration time evaluated after a delay of 8 hours, n= 7 WT and 5 PRKO, * p=0.029 (t=2.499, df=11), student’s t-test. **(D)** The percent novel object exploration time evaluated after one-hour delay, n= 7 WT and 10 PRKO, * p=0.0006 (t=4.291, df=15), student’s t-test. **(E)** Average and standard deviation of the total exploration time in the WT and PRKO mice during familiarization and testing phases, n= 7 WT and 10 PRKO. **(F)** Movement of representative WT and PRKO mice in the open-field arena. **(G)** Average and standard deviation of the total distance traveled by 7 WT and 10 PRKO mice. **(H)** The percentage novel object exploration of the vehicle- and RU-486-treated animals, n=9 vehicle-treated and 7 RU-486-treated, * p=0.0165 (t=2.724, df=14) student’s t-test. The animals were treated daily with vehicle (20% β-hydroxycyclodextrin daily subcutaneous, sc) and RU-486 (10 mg/kg, sc, daily) for a week. There was a one-hour delay between the familiarization and testing phases. **(I)** Average and standard deviation of the total exploration time in the WT and PRKO mice during familiarization and testing phases, n= 9 vehicle, and 7 RU-486-treated animals.

We assessed whether the distinction in the object exploration or movements around the arena could account for the observed differences between the WT and PRKO mice. But, two genotypes spent the same amount of time exploring objects during the familiarization and testing phases (Fig. 5E) and traveled comparable distances (Fig. 5F, 5G). Thus, the deficits in PRKO mice were not due to altered object exploration or locomotion.

Progesterone levels could have contributed to the observed differences in the novel object recognition in the PRKO and WT mice; therefore, we measured the progesterone levels in the combined EC and hippocampal tissue lysates from the PRKO and WT mice using an ELISA assay. The progesterone levels in these tissues were similar, WT males: 20.5 ± 5.7 pg/mg, n=6, and PRKO males: 21.1 ± 9.7 pg/mg, n=5. These levels were similar to the progesterone levels in females in the estrus stage of the cycle (16 ± 6 pg/mg, n=5).

### The pharmacological blockade of PRs also affected novel object recognition

An absence of PRs throughout the development of the PRKO mice may have contributed to the observed deficits in adult mice. We tested whether blocking the PR activity for a short period in adult mice also impacted object recognition. We treated adult male mice with the PR antagonist RU-486 daily for a week and evaluated its effect on novel object recognition. Like the PRKO mice, the RU-486-treated animals could also not distinguish the novel from the familiar object (Fig. 5H). The vehicle- and RU-486-treated animals spent comparable time exploring the two objects during the familiarization and testing phases (Fig. 5I), indicating that the suppression of PR signaling did not affect exploration. Despite the similarity in locomotor activity, the lack of specificity in the exploration of the novel object during the testing phase indicates that the pharmacological blockade of PRs in adulthood interferes with short-term memory processes necessary to detect environmental changes such as that presented during the novelty test.

### Novel object recognition was also affected in the female PRKO mice

Because progesterone is a canonical female reproductive hormone, we also determined whether the role of PRs in the regulation of novel object recognition uncovered in males was also critical in females. The PRKO mice experience irregular estrous cycles, so the animals were ovariectomized to control hormone levels. We tested them 12-14 days later. The overiectomized PRKO and WT female animals did not differ in object recognition evaluated after an hour of delay (Fig. 6A). This was expected since these animals’ circulating and brain progesterone levels were negligible (data not shown). Progesterone treatment improved novel object recognition of the WT but not the PRKO (overiectomized) animals (Fig. 6B, overall p=0.042, ANOVA, p=0.046 šídák’s post-hoc multiple comparisons test WT V vs WT P).

**Figure 6:**
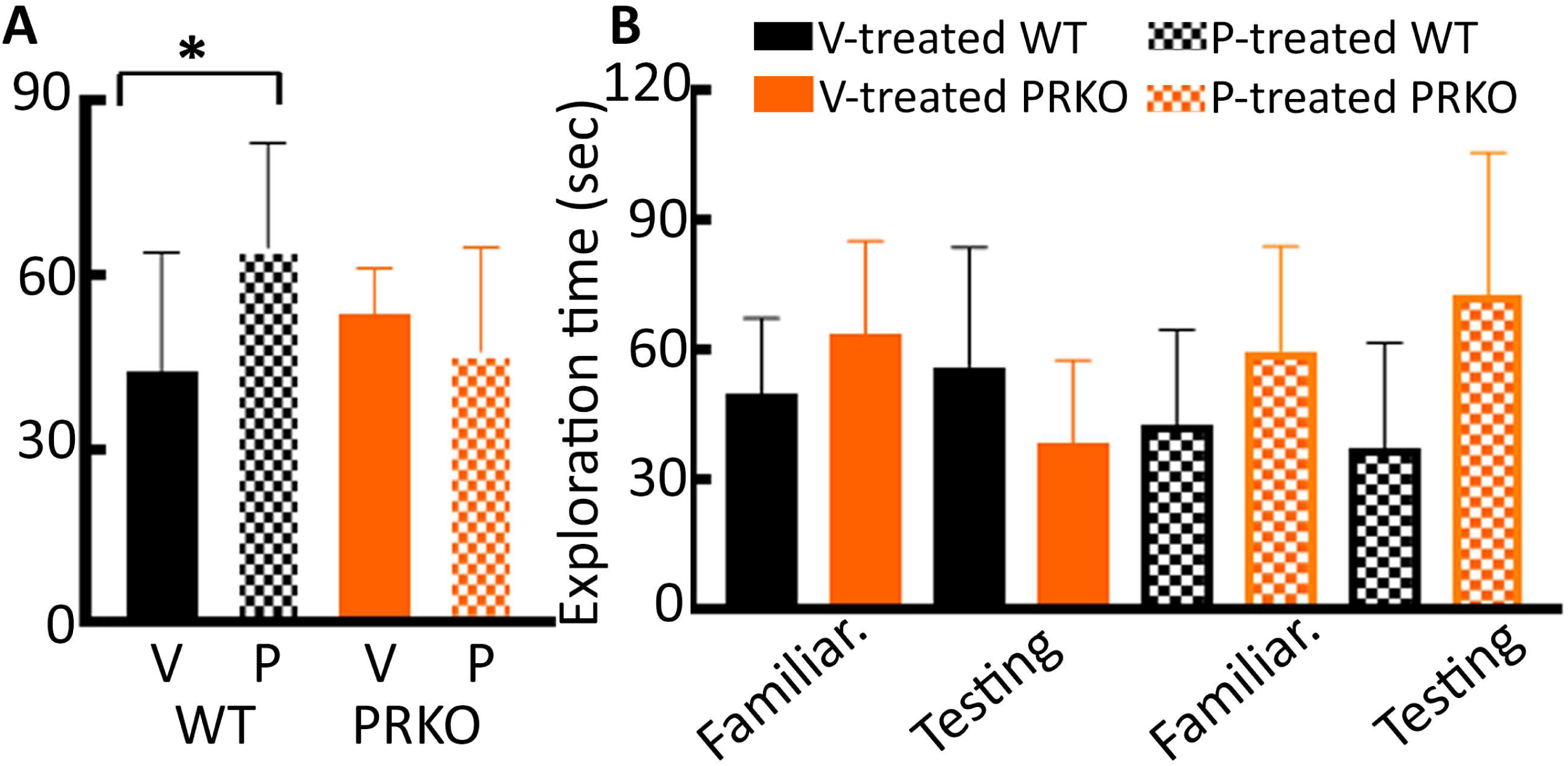
The effect of PR deletion on novel object recognition in female mice. **(A)** The percent novel object exploration time in the ovariectomized WT and PRKO female mice treated with vehicle or progesterone (10 mg/kg, sc) for 2 consecutive days. The familiarization and testing were performed 24 hrs after the last progesterone/vehicle injections, and there was an hour delay between the familiarization and testing, n= 9 vehicle-treated and 10 progesterone-treated WT animals and 6 vehicle-treated and 7 progesterone-treated PRKO animals, * p=0.037 ANOVA with šídák’s multiple comparisons test between V and P-treated WT animals. **(B)** The total object exploration time during the familiarization and testing phases. The number of replicates is the same as in panel A.

The PR deletion or progesterone treatment did not affect the time of exploration during the familiarization or testing phases (Fig. 6B). Thus, the PR signaling was also critical for regulating novel object recognition in females.

### Male PRKO mice could also not recognize object place change

Episodic memories comprise “what”, and “where,” components (Tulving, 1972; Winters et al., 2008), and we found an impact of PR deletion or their pharmacological blockade on the “what” component. We evaluated whether PRs also regulated the “where” element, i.e. recognizing the object’s place (Fig. 7A). The male PRKO mice also differed from the WT mice in recognizing the object’s place change. The WT mice explored the displaced object more (Fig. 7B, 7C), but not the PRKO mice (Fig. 7B, 7C, n=8 each, p=0.016, t=2.728, df=14, student’s t-test). Together, these studies revealed a critical role of PR signaling in regulating spatial memories in male and female mice. In the subsequent analyses, we evaluated whether PR signaling also regulated other forms of memories.

**Figure 7:**
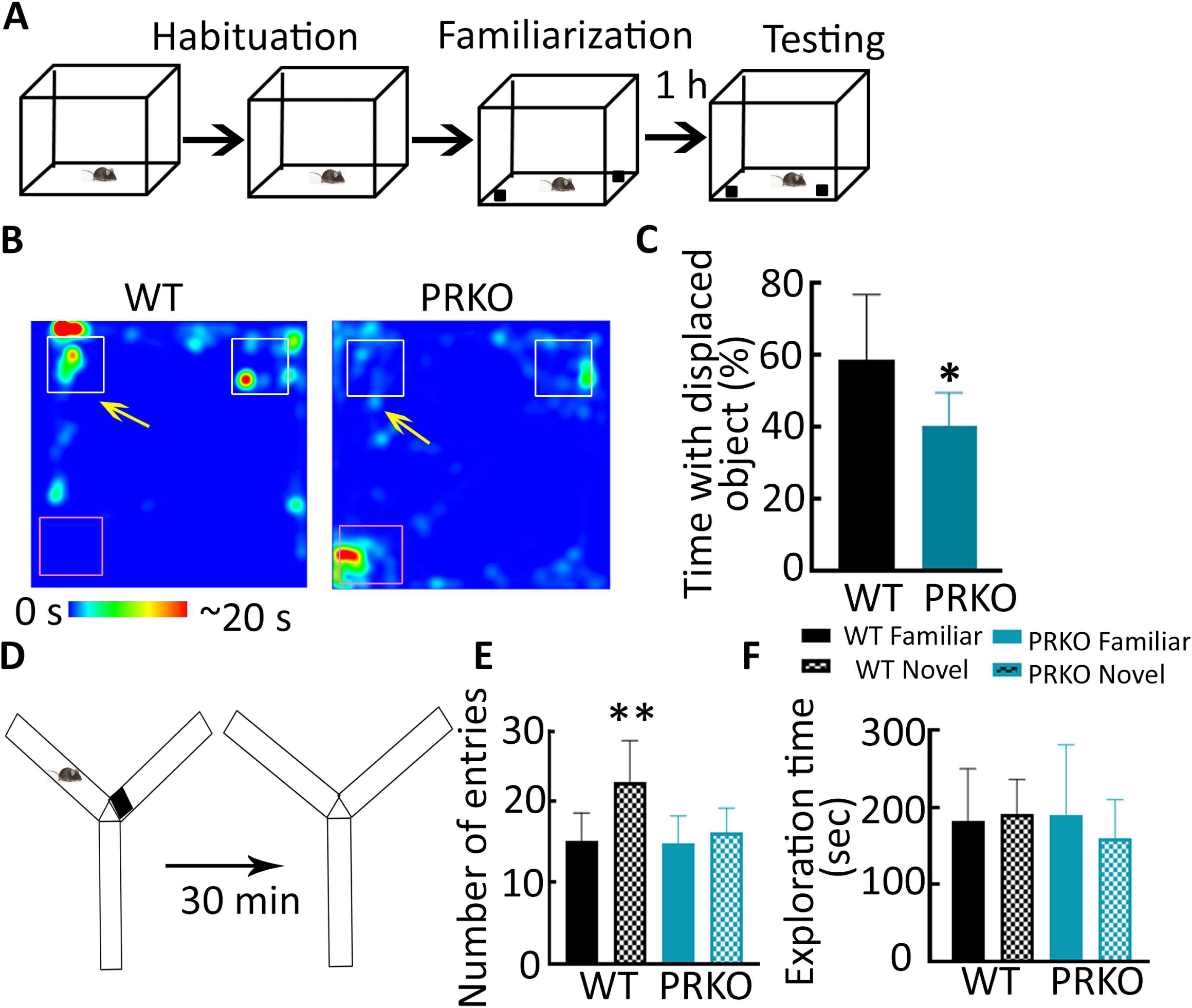
The effect of PR deletion on object place recognition and Y maze forced-alternation. **(A)** The schematic showing object place recognition testing. **(B)** Heat map from representative WT and PRKO mice during the testing phase. The arrows mark the displaced object; their original position is marked by black squares. The testing was performed after an hour of delay. **(C)** The percent time spent in exploring the displaced object, n=8 each, * p=0.016 (t=2.728, df=14) student’s t-test. **(D)** A schematic of the Y maze forced-alternation test. **(E)** The number of entries in the novel arm by PRKO and WT males in a Y maze forced-alternation task, n= 6 each, overall p= 0.0074, * p=0.0031, ANOVA with Sidák’s multiple comparison WT familiar vs WT novel. **(F)** The time spent exploring the novel and familiar arms. The number of replicates is the same as in E.

### EC cortex-specific deletion of PRs was sufficient to impair novel object recognition

The PRKO mice had impaired novel object recognition; however, these mice lacked the PRs in the entire brain. Furthermore, the PRs were also absent throughout the development, which may have exerted additional effects. We evaluated whether the entorhinal cortex-specific deletion of PRs in adult mice affected object recognition. A GFP-tagged Cre recombinase was expressed in the EC cortex of adult males using AAV vectors, which transduced 70-80% of the entorhinal cortical neurons (Fig. 8A) and reduced the PR mRNA expression (45 ± 16%, n=4). After confirming the EC cortex-specific deletion of PRs, we tested the Cre-expressing animals in the novel object recognition task. The Cre-expressing mice did not show greater interest in the novel object (Fig. 8B), as seen with the mice expressing only GFP in the EC cortical neurons (n=8 each, p=0.0204, t=2.613, df=14, student’s t-test). Thus, blocking the progesterone-PR signaling in the EC cortical neurons was sufficient to impact object recognition.

**Figure 8:**
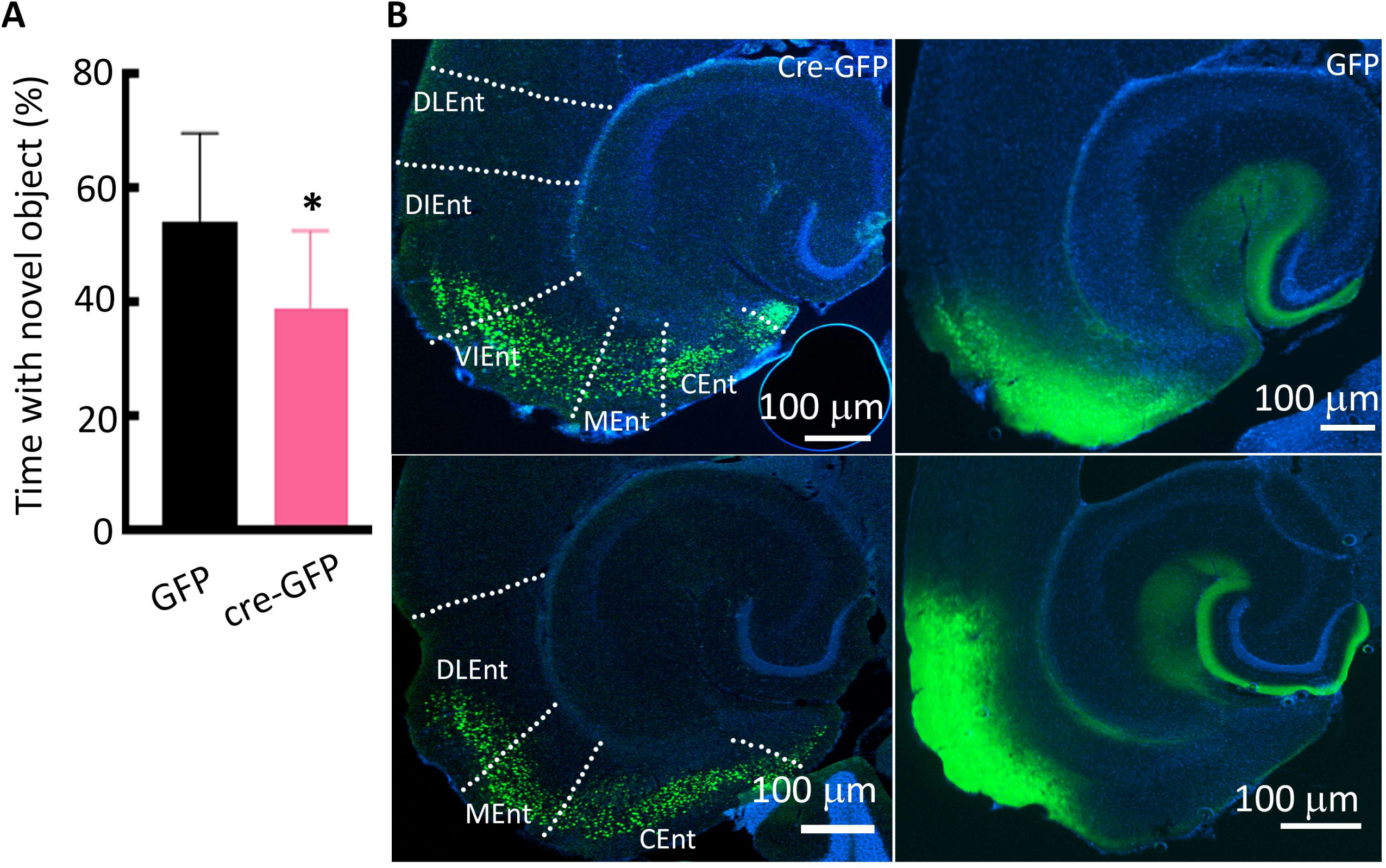
Entorhinal cortex-specific deletion of PRs and novel object recognition. **(A)** Percent novel object exploration in the animals injected with AAVs to express CamKII driven Cre-GFP or GFP alone, n=8 each, *0.0204 (t=2.613, df=14) student’s t-test. **(B)** Images from representative animals injected with AAVs to express CamKII-driven Cre-GFP or GFP alone. Sections from the dorsal and ventral EC cortex are shown, the green fluorescence corresponds to Cre-GFP or GFP expression and DAPI (blue) was used as a counterstain.

### Spatial reference memory was affected in PRKO mice

We confirmed that PRs regulate spatial memories using a Y-maze forced-alternation task (Kraeuter et al., 2019). This task also does not require extensive training or subject animals to stress. One arm was blocked during the initial exposure; mice were placed in the home cage for thirty minutes, then brought to the maze with both arms open (Fig. 7D). The WT mice entered the novel arm more often (Fig. 7E), the PRKO mice, in contrast, visited the familiar and novel arms equally (Fig. 7E, n= 6 each, overall p=0.0074 ANOVA, p=0.0031 Sidák’s multiple comparisons WT novel vs WT familiar). The time spent in the two arms did not differ in either genotype (Fig. 7F). Thus, we confirmed that PR deletion impacts spatial reference memory.

### Contextual fear conditioning was not affected in the PRKO mice

In contrast to spatial memories, implicit memories include procedural memories or those driven by stimulus or priming and do not involve a conscious recall of the events. We evaluated fear conditioning in WT and PRKO male mice to understand whether PR signaling also regulates implicit memories (Fig. 9A). Animals’ immobility and freezing increased after each foot shock on the day of conditioning, and this was similar between the PRKO and WT mice (Fig. 9B). Freezing during the context recall testing performed 24 hours later was identical in the two groups (Fig. 9C, p=0.47, t=0.7512, df=9, student’s t-test). Thus, PR deletion did not impact associative memory.

**Figure 9:**
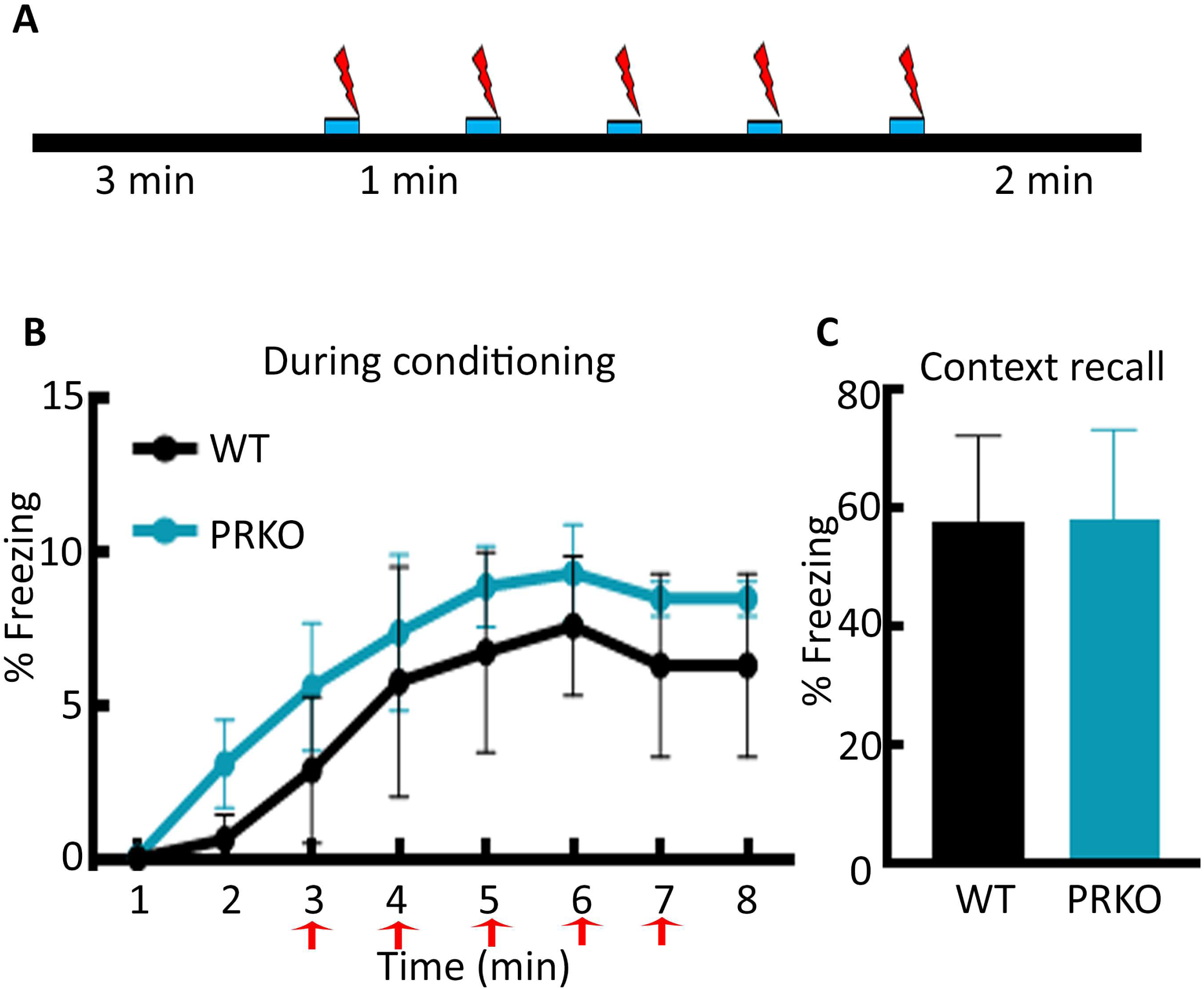
The cued-contextual fear condition in PRKO male mice. **(A)** A schematic showing fear conditioning and recall. Following an initial 3 min of chamber exploration, the animals received five 2-sec footshocks that were contiguous with a 20-sec tone. The animals were removed from the chamber two min after the last footshock. **(B)** Percent freezing in the WT and PRKO male mice following the five footshocks, n= 6 WT and 7 PRKO. **(C)** Percent freezing during the context recall was performed 24 hours later. The number of replicates is the same as in E.

## DISCUSSION

We demonstrate that principal neurons in the EC-hippocampal circuit express PRs, ablating them diminishes spatial, declarative memories. The following observations support this conclusion. First, PRs were expressed primarily in principal neurons within the EC and hippocampus of male mice. Activation of PRs with segesterone, increased the number of active neurons in the DGC, CA1, subicular, and EC. In contrast, disruption of PR signaling either with RU 486 or genetic ablation impaired spatial memories, short and long term but not fear conditioning. The PR-regulated spatial memory may offer reproductive advantages, helping animals find mates and food.

We found that PRs regulate spatial memory. The lateral and medial EC relay non-spatial and spatial information to the hippocampus, which integrates these inputs (Witter et al., 2000; Manns and Eichenbaum, 2006). The hippocampal CA1 neurons (place cells) have a tight coupling of firing to their fields (O’Keefe John 1978). The medial EC neurons (grid cells and border cells), particularly those in the superficial layers, also have strong space firing fields (Quirk et al., 1992; Fyhn et al., 2004; Hargreaves et al., 2005; Moser et al., 2014). The medial EC neurons receive sensory inputs from visual and parietal cortices through the post-rhinal cortex (Burwell et al., 2004; Kerr et al., 2007). In contrast to the medial EC, the spatial firing selectivity of lateral EC neurons is weak (Kerr et al., 2007), and these neurons are proposed to function as object-responsive cells and object trace cells (Tsao et al., 2013). Lesioning of the lateral EC affects recognition of object, its place, and context (Van Cauter et al., 2013; Wilson et al., 2013a). Both the medial and lateral EC neurons project to the DGCs via the perforant pathway, and lesions of the dentate gyrus also affect the recognition of objects’ place (Kesner et al., 2015). This indicates that impairing EC-DG function impacts spatial memory. The potentiation of glutamatergic synaptic transmission termed long-term potentiation (LTP) is critical for memory formation; its blockade or saturation impairs cognitive processes (Moser et al., 1998; Whitlock et al., 2006; Nicoll, 2017; Naik et al., 2020). The strong PR expression in the EC and hippocampi seen here taken together with the role of PRs in regulating AMPA receptor-mediated transmission of these neurons found in earlier studies (Joshi et al., 2018; Shiono et al., 2021), raises the possibility that PR activity may regulate LTP these neurons. Indeed, affected EC-DG communication is seen during aging and in Alzheimer’s disease (AD), conditions associated with cognitive decline. A loss of perforant path-DGC synapses in the outer molecular layer is seen in patients with mild cognitive impairment and early AD (Scheff et al., 2006). In contrast, aging modifies the lateral perforant path-DGC synapses plasticity (Froc et al., 2003). A putative loss of PRs or PR-expressing neurons combined with declining progesterone levels could affect the normal transmission between EC-DG-CA1 neurons, which is critical for spatial memory.

Prior studies have reported PR expression in the limbic regions, including the amygdala and hippocampus of the fetal brain (Quadros et al., 2007; Newell et al., 2018) and the adult female brain (Guerra-Araiza et al., 2000; Guerra-Araiza et al., 2003; Waters et al., 2008; Mitterling et al., 2010; Shiono et al., 2021). In contrast, extrahypothalamic PR expression in adult male brains is not characterized, although prior studies have suggested that PRs are expressed in the hippocampi (Guerra-Araiza et al., 2001; Mitterling et al., 2010). We found that PR expression in the EC-hippocampal network was comparable between adult males and females. Furthermore, the PR-expressing cells in these regions also expressed CamKII, a marker of glutamatergic neurons, but not the markers of interneurons, PV and Som peptides, indicating their excitatory nature. Although we evaluated the colocalization of tdTomato with neuropeptides marking only two interneuron types, the high degree of colocalization with CamKII and a near complete lack of colocalization with interneuronal markers suggests that PR are not expressed in interneurons. The findings in dentate hilus, rich in interneurons, also support this as the PR-expressing cells expressed GluA2 subunit, a marker for hilar mossy cells, but not the interneuronal marker peptides parvalbumin and somatostatin. A prior study in neonatal animals did not find PR immunoreactivity overlap with that of GAD67 in the dentate gyrus (Newell et al., 2018).

Sex differences exist for various cognitive tasks; women perform better in episodic memory tasks, whereas men have an advantage in visuospatial tasks (Miller and Halpern, 2014; Asperholm et al., 2019). However, the role of progesterone-PR signaling is shared in males and females, which is distinct from the often opposing actions of PR signaling in the mating behavior of males and females (Wagner, 2006). Correct identification of an object/individual depends on properly integrated sensory cues. PRs may regulate the generation and/or integration of sensory cues, and their deletion could affect these functions. Although not typically thought of as reproductive behavior, spatial and recognition memories are also critical for successful reproduction. Since, the home range of males (462 m^2^) is double that of females (214 m^2^) (Mikesic and Drickamer, 1992), spatial memory and object identification could be more critical in them than in females. Sexual selection favors mating of individuals with better traits; a better spatial memory could help territory marking by males, as seen in lekking hummingbirds (Araya-Salas et al., 2018). Females also appear to prefer males with a better spatial ability in meadow voles (Spritzer et al., 2005). A better spatial memory could favor sexual selection (larger size) by increasing foraging avenues through a better memory of food source locations. An earlier study found that PRKO males are less aggressive (Schneider et al., 2003), which raises the possibility that territory marking and defense may be affected. Hence, although reproduction per se does not seem to be affected by PR deletion in males in laboratory conditions, absence of this signaling could be disadvantageous for reproduction in males in real world.

PR expression in the Cajal-Ritzius cells of the DG of developing mice is proposed to regulate the perforant path innervation of the molecular layer, inhibiting PRs with RU486 during the first week of life affects episodic-like memory in adult males (Newell et al., 2018). The lack of PRs throughout the development of PRKO mice could have affected the EC-DG synapses. However, this effect was not exclusively due to the developmental absence of PRs, since, RU486 treatment or EC-specific PR deletion in adulthood also impacted novel object recognition. Thus, impairing the EC-DG network activity may be sufficient to alter episodic-like memory formation.

We have previously found that PR activation increases the excitability of entorhinal cortical neurons in adult females (Shiono et al., 2021). A similar effect of PR activation in male mice predicted that treatment with PR agonist segesterone would increase the number of active neurons in the entorhinal cortical-hippocampal network. Indeed, our experiments using TRAP mice in which activity-dependent promoter cFos drives the expression of tdTomato reporter protein revealed an increased number of active neurons in the hippocampus and entorhinal cortex. Surprisingly, however, the TRAPed neurons in the lateral entorhinal cortex were spread over all the layers in contrast to the predominant expression of PRs in the superficial layers, which indicates that PR agonist treatment likely activated PR-expressing neurons and their target neurons.

In conclusion, the excitatory neurons of EC and hippocampi express PRs; exciting these receptors leads to neuronal activation. PRs facilitate spatial memories.

## Acknowledgments

This study was supported by the National Institutes of Health (NIH) grants R01 NS 110863 to SJ. The content is solely the responsibility of the authors and does not necessarily represent the official views of the National Institutes of Health. We thank Drs Howard P Goodkin and Anastasia Brodovaskaya, and John Williamson for critical comments, and Dr. Aijaz Naik for help with setting up the apparatus.

